# Weighted trait-abundance early warning signals better predict population collapse

**DOI:** 10.1101/282087

**Authors:** Christopher F. Clements, Martijn van de Pol, Arpat Ozgul

**Affiliations:** Department of Evolutionary Biology and Environmental Studies, University of Zurich, Zurich 8057, Switzerland; School of BioSciences, University of Melbourne, Melbourne 3010, Australia; Department of Ecology & Evolution, Research School of Biology, Australian National University, Canberra 2601, Australia; Department of Animal Ecology, Netherlands Institute of Ecology (NIOO-KNAW), Wageningen 6708PB, the Netherlands

**Keywords:** Body size, extinction, population dynamics, tipping points, trait dynamics

## Abstract

Predicting population collapse in the face of unprecedented anthropogenic pressures is a key challenge in conservation. Abundance-based early warning signals have been suggested as a possible solution to this problem; however, they are known to be susceptible to the spatial and temporal subsampling ubiquitous to abundance estimates of wild population. Recent work has shown that composite early warning methods that take into account changes in fitness-related phenotypic traits - such as body size - alongside traditional abundance-based signals are better able to predict collapse, as trait dynamic estimates are less susceptible to sampling protocols. However, these previously developed composite early warning methods weighted the relative contribution of abundance and trait dynamics evenly. Here we present an extension to this work where the relative importance of different data types can be weighted in line with the quality of available data. Using data from a small-scale experimental system we demonstrate that weighted indicators can improve the accuracy of composite early warning signals by >60%. Our work shows that non-uniform weighting can increase the likelihood of correctly detecting a true positive early warning signal in wild populations, with direct relevance for conservation management.

## Introduction

Statistical early warning signals (EWSs) calculated from abundance time series data have been suggested as a possible method for predicting approaching population collapses and regime shifts (Drake & Griffen, 2010; Carpenter et al., 2011; Dakos et al., 2012; Kéfi et al., 2013). However, abundance-based early warning signals are known to be susceptible to the spatial and temporal subsampling ubiquitous to wild population abundance estimates (Clements et al., 2015), and have been criticized for not reliably predicting significant declines in natural populations (Burthe et al., 2016). Recent work has sought to resolve these issues by incorporating data on the dynamics of fitness-related phenotypic traits alongside abundance data (Clements & Ozgul, 2018). Traits such as body size are highly responsive to environmental perturbations and changes in the dynamics of these traits often precede demographic responses to deteriorating environments (Anderson et al., 2008; Ozgul et al., 2014; Clements & Ozgul, 2016a). Incorporating information on the shift in the body-size distribution of a population can not only provide an additional measure of stability (Anderson et al., 2008), but has the potential to improve the predictive accuracy of EWS as trait dynamic estimates may be less susceptible to sampling protocols than population abundance estimates are when the distribution of ages and sexes is assumed to be random (spatial partitioning between ages or sexes may affect this) (Clements et al., 2015, 2017; Clements & Ozgul, 2016a). Previous work has shown composite early warning metrics that include data on both abundance and trait dynamics better predict population collapse than those that incorporate abundance-only or trait-only data (Clements & Ozgul, 2016a).

Recently developed trait-abundance composite early warning indicators have been based upon the method proposed by Drake & Griffen (2010), whereby multiple statistical signals are normalized and then summed to create a single composite signal. Clements & Ozgul (2016a) used this approach to incorporate shifts in mean body size and variance in body size along with concurrent changes in the statistical properties of an abundance time series, and demonstrated that such an approach can significantly improve the reliability of early warning signals in both experimental (Clements & Ozgul, 2016a) and natural (Clements et al., 2017) populations. However, in this method the relative importance of abundance versus trait data in the composite indicators was weighted evenly. Given the known issues with abundance data, a logical extension to this method is to non-evenly weight the relatively importance of abundance and trait data in the composite indicators.

Non-uniform weighting of model parameters has a history of use in conservation biology, particularly in determining optimal management strategies to maximize outputs from limited resources (Joseph, Maloney & Possingham, 2009). For example habitat conservation may be prioritized based on the suitability of the habitat for certain species, with such weightings often being determined by expert opinion (Lehtomäki et al., 2009). Such approaches have also been used to assess trade-offs, for example between conservation and carbon sequestration (Thomas et al., 2013). As well as expert opinion, weighting may be based on more quantitative measures of data quality; for example by the frequency of sampling of a population to estimate abundances, or the percentage of a habitat sampled when counting individuals, both of which have been shown to affect the reliability of early warning signals (Clements et al., 2015). However, practitioners must first discern if non-uniform weightings convey an advantage before implementing such an approach for wild populations.

Here we assess whether non-uniform weightings improve the predictive ability of composite EWS of population collapse using data from an experimental protozoa study. We take the most reliable composite early warning metric (as identified by Clements & Ozgul (2016a)), and alter the relative weighting of the abundance and trait data when calculating whether an early warning signal is present or not. We then reanalyze the data from an experimental protist microcosm system, presented in Clements & Ozgul (2016a), and show that alternate weightings can improve the predictive ability of composite EWS by decreasing the frequency of false positive signals, and increasing the frequency of true positive signals.

## Methods

### Experimental Data

Data on the population dynamics and body-size (width, μm – a proxy for mass) of individuals of a predatory ciliate protozoa (*Didinium nasutum*) feeding on a bactiverous ciliate protozoa (*Paramecium caudatum*) were collected over a 47-day period (Fig. 1). Populations of *D. nasutum* were subjected to four different treatments (15 replicates per treatment), where the number of *P. caudatum* fed to each population per day was manipulated. In one treatment (“Constant”) populations of *D. nasutum* were fed 300 *P. caudatum* per day for the 47 days of the experiment, whilst in the other three treatments the number of *P. caudatum* declined through time at three different rates (“Slow”, “Medium”, “Fast”) driving the populations of *D. nasutum* to extinction at varying points in time, and with varying population dynamics prior to extinction (Fig. 1). None of the populations in the Constant treatment went extinct. For each population the time at which it passed through a tipping point, if at all, was calculated (as in Drake & Griffen, 2010), and early warning signals were then calculated prior to the occurrence of each of these tipping points. Because of the size of the microcosms it was impractical to count every individual of a population, hence a subsample was taken (10% of the habitat, a volume that allowed all individuals to be easily counted with close to zero error) and we assumed that the total number of individuals in each microcosm was reflected by the abundance in the subsample. Whilst this undoubtedly introduced some minor error into the abundance estimates, EWSs were still detectable using this uncorrected subsample data (Clements & Ozgul, 2016a). We believe that this uncertainty in abundances is very representative of the ubiquitous spatial subsampling associated with the monitoring of all wild populations, and hence applying such methods to this data is a reliable reflection of the challenges of applying them to real world population dynamics. For full details of the experimental design and protocols see Clements & Ozgul (2016a).

**Figure 1.**
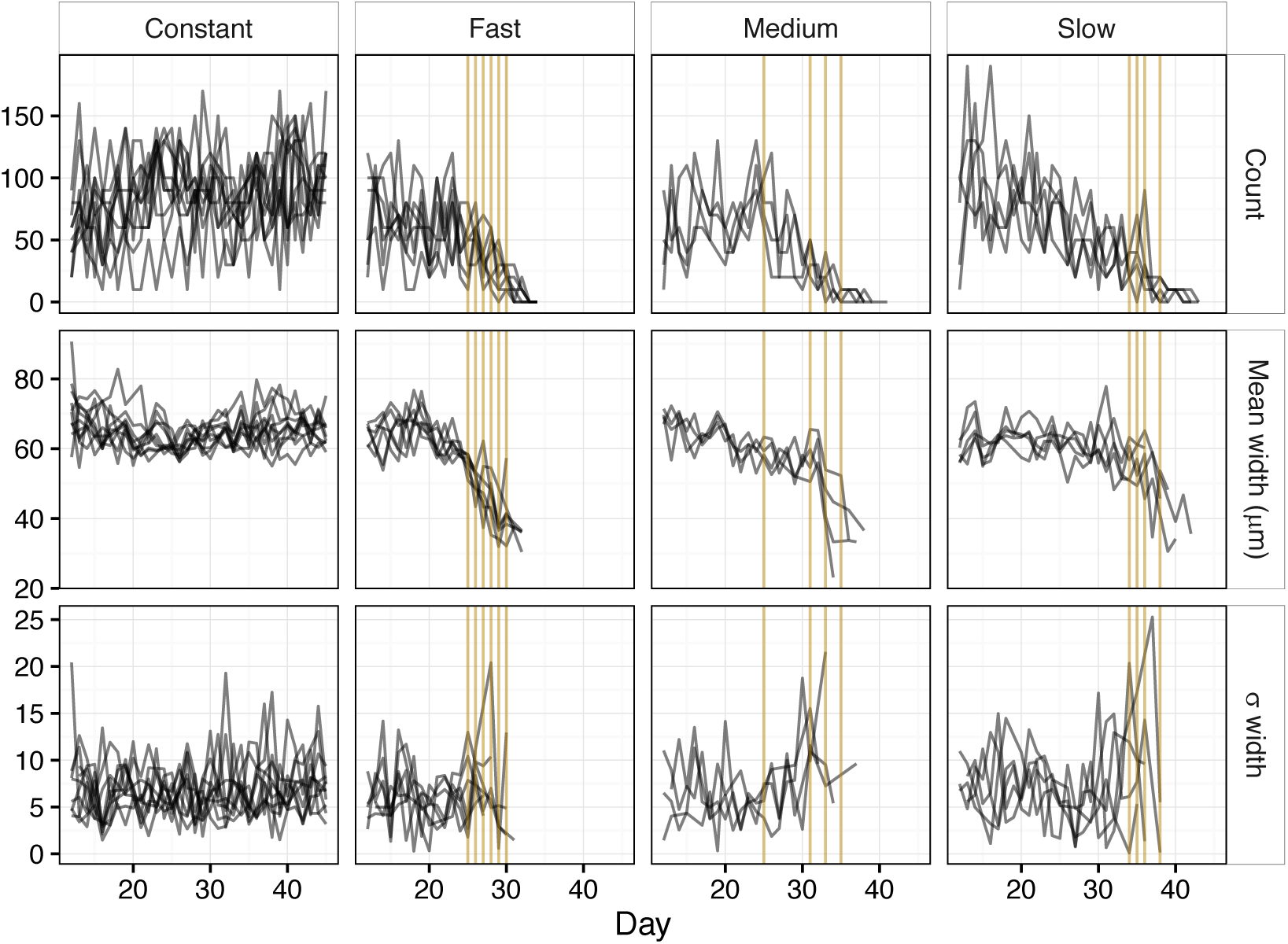
Black lines describe the population and body size dynamics of individual populations of *Didinium nasutum* subjected to four different experimental treatments (constant, fast, medium, and slow rates of decline in prey availability). Data from day 0 to 12 were removed to minimize the effects of transitory dynamics. Each vertical gold line indicates an inferred tipping points for a collapsing population.

### Early warning signals

Previous work has identified a composite index comprised of the coefficient of variation of the abundance time series (*cv*), shifts in mean body size of the individuals in the population (*size*), and shifts in the standard deviation of mean body size (*sd.size*) as producing the most reliable estimates of whether a population was at risk of collapse in these experimental data (Clements & Ozgul, 2016a). Here we test this composite index by systematically altering the weighting of these three competent parts as a proof of concept of non-uniform weighting increasing the predictive accuracy of the composite metric.

Here we implement the approach developed by Clements & Ozgul (2016a). Each of the three leading indicators (*cv, size, sd.size*) was calculated at each day observations were made, and for each experimental population independently. Each leading indicator was then normalized by subtracting the long-run mean of that indicator from the value of that indicator at each time point, and dividing it by the long run standard deviation (Drake & Griffen, 2010; Clements & Ozgul, 2016a) (Supplementary Information). The value of the composite early warning signal was then calculated by summing the value of each leading indicator (*cv, size, sd.size*) at each time point. Previous work has suggested an EWS could be considered present when the value of this composite EWS exceeds its running mean by either 1 or 2σ (Drake & Griffen, 2010). Recent evidence suggesting a 2σ threshold provides more reliable results (Clements & Ozgul, 2016a) and consequently here we consider a signal present at a 2σ threshold.

The weighting of each of the three leading indicators was altered by multiplying the normalized value of each metric prior to summing them together to calculate the composite EWS. Each leading indicator was weighted from 1 to 10, with every combination of weightings tested (e.g. *cv*_*w*_=1:*size*_*w*_=2:*size.sd_w_*=5, *cv*_*w*_=8:*size*_*w*_=4:*size.sd_w_*=1). The performance of each weighting was assessed by using a “normalized metric score” (Clements & Ozgul, 2016a), calculated by subtracting the proportion of false positives (EWS present in data from the constant treatment) from the proportion of true positives (EWS present in data from the slow, medium, and fast treatments). The highest scoring weighting for each of the slow, medium, and fast treatments was compared to uniform weighting in each of these treatments (Fig. 2a). The best metric when data from all three treatments were grouped together was calculated as the weighting with the highest normalized metric score, and the minimum difference in normalized metric scores between treatments (Fig. 2b). This gave an indication as to the weighting that was most robust to different rates of environmental change, and thus potentially most widely applicable to different scenarios.

**Figure 2.**
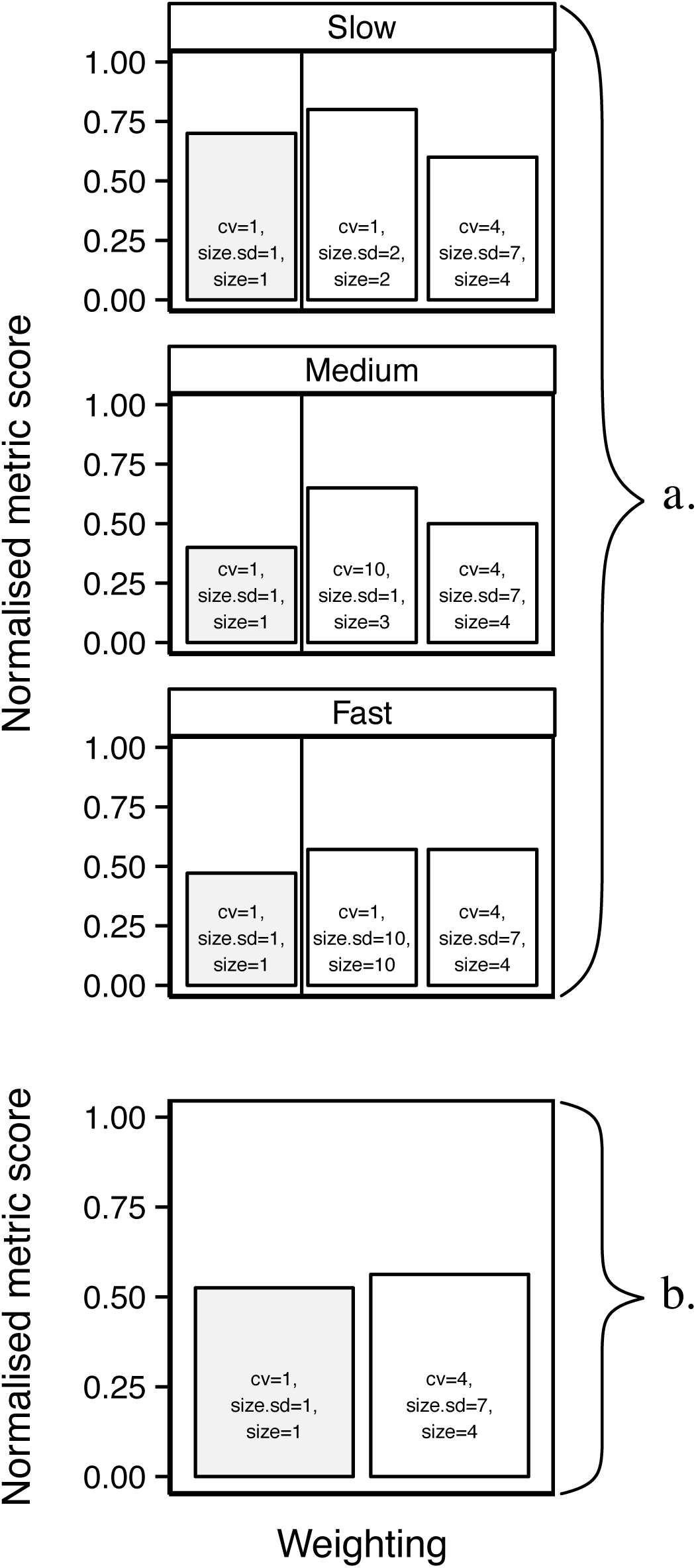
(a) The highest scoring weighting across each of the experimental treatments compared to even weighting and the best weighting when data from all treatments were combined, and (b) the weighting with the highest normalized metric score across all three treatments, and the lowest difference in normalized metric score amongst treatments.

All analyses were carried out using the statistical software R (R Development Core Team, 2016), and the code to implement weighted trait-abundance early warning signals is available as supplementary information.

## Results

### Experimental data

Non-uniform weighting of the importance of abundance and tra it data in early warning indicators can improve the reliability of these methods in predicting population collapses (Fig. 2). The largest improvement (62.5%) was seen when data from the Medium treatment was analyzed, possibly because uniform weighting performed relatively poorly (Fig. 2a). The highest achieved normalized metric score was 0.8 (in the slow treatment), suggesting very high numbers of true positive EWS, and low numbers of false positive EWS (Fig. 2a).

When data from three deteriorating treatments was grouped together the weighting that produced the greatest improvement in predictive accuracy weighted the relative importance of *cv, size.sd, size* as 4:7:4, although the improvement over uniform weighting was not large (Fig. 2b), suggesting that the how fast the pressure on the system changes (known as the rate of forcing (Clements & Ozgul, 2016b)) may be an important factor in determining not only the correct weighting to apply, but also our ability to reliably predict population declines. To highlight this, the 4:7:4 weighting performed as well as the best weighting in the fast treatment, average in the medium treatment, and worse than both the uniform and best weighting in the slow treatment (Fig. 2a).

## Discussion

Predicting population collapse is a key but challenging goal in conservation biology. Because previously developed EWS that take into account both trait and abundance data are non-system specific, and thus widely applicable, they may be of particular interest. Here we analyze data from a small-scale experimental system and show that non-uniform weighting can improve the reliability and strength of trait-abundance early warning signals, but that the strength of this improvement is not uniform across different rates of environmental change.

Previous work in small-scale experimental systems has identified a composite metric of *cv, size*, and *size.sd* as the most reliable predictor of population collapse in experimental microcosm populations. This method provides improved reliability over methods that are based on either abundance-only or size-only data; however, the method still produces false positive and false negative signals in some populations. Because of the known susceptibility of abundance-based early warning signals to poor quality data (Clements et al., 2015), non-uniformly weighting the components of composite metrics provides an obvious extension to this previous work.

Here we demonstrate, using the same experimental data with which the original trait-abundance method was developed, that a weighting of 4:7:4 (*cv:size.sd:size*) provides the greatest overall improvement across all three treatments, with the minimum between-treatment variation in this result (Fig. 2b). Resilience to treatment variation in performance in the experiment is important, as it maximizes the reliability of applying such methods to systems where the rate of change remains unknown. However, the among-treatment variation in the potential advantages of non-uniform weighting should not be ignored, as with weightings other than 4:7:4 there were significantly higher normalized metric scores in the medium and slow treatments (Fig 3a). Such a result is likely to be driven by the rate of forcing, known to potentially alter the detectability of EWSs (Clements & Ozgul, 2016b), of the system altering the rates of change of the mean and σ body size of individuals. For example, mean body size rapidly declines in the fast treatment (Fig. 1), whilst in the medium treatment body size decline is more gradual and a weighting towards the coefficient of variation of abundance, rather than towards body size, improves predictive accuracy (Fig. 2). These results suggest that the rate of forcing a system can alter the weighting that produces the most reliable predictions of an approaching population collapse.

Generalizing such a result to real-world systems may be problematic, as we cannot assume that the population and trait dynamics of the microcosm system analyzed here are truly representative of all real-world population collapses. Ideally one would select the weighting based on an estimate of the reliability of the available abundance or trait data, and possibly based on the rate of forcing of the system, although doing so is likely to be non-trivial. If, for example, available abundance data are known to be estimated from a survey conducted on a small proportion of the known range of a species, or are temporally limited, it may be prudent to calculate the presence of early warning signals with a bias in favor of trait-based data. A less quantitative option would be to weight metrics based on expert opinion of the reliability of the available data. However, whilst criticism has been leveled at the use of expert opinion in conservation management, it has been shown to be useful if approached with caution (Johnson & Gillingham, 2004; Martin et al., 2005, 2012). Similar caution must be applied to the non-uniform weighting of trait and abundance data in the models presented here, however the significant improvements in the predictive accuracy of these approaches when weighting is non-uniform mean that canvasing expert opinion may be a relatively simple and cost-effective solution to improve predictive accuracy.

In conclusion, we demonstrate the possible advantages of non-uniform weighting in an early warning signals framework. This work provides a first step to improving the reliability of recently proposed abundance-trait methods (Clements & Ozgul, 2016a), and may be used to negate some of the known issues that affect abundance-based EWSs (Clements et al., 2015). Future work may seek to make more concrete recommendations on weightings based on qualitative measures such as expert opinion, or more quantitative measures such as measures of data quality, the known level of threat to a species or a population, the trophic level of the species, or its connectedness in a network. One option to tackle this is to use complex size-structured community models, such as those commonly used in fisheries (Blanchard et al., 2012; Scott, Blanchard & Andersen, 2014), to simulate shifts in the trait dynamics and abundances of multiple interacting species, allowing alternative weightings of data from various trophic levels to be tested on communities where collapse can be invoked by, for example, overfishing or changing climatic variables.

## Acknowledgements

This work was made possible by an SNSF Post-doctoral fellowship (P300PA_174359/1) awarded to C.C. and an ERC Starting grant (#337785) to A.O. M.vd.P. was supported by a Future fellowship from the Australian Research Council (FT12010020).

